# Isolation and Genomic Characterization of *Myxococcus faecalis* Strains from Mangroves in Southeastern Brazil

**DOI:** 10.64898/2026.04.28.721309

**Authors:** Renan S. Oliveira, Yann F. Lin, Paula C. Jimenez

## Abstract

*Myxococcus faecalis* was recently described from human fecal isolates, although subsequent evidence indicates an environmental distribution for this lineage. Here, we report the isolation and genomic characterization of two *M. faecalis* strains (BRX-014 and BRX-032) recovered from mangrove ecosystems along the southeastern coast of Brazil, representing the first record of the species in a marine-coastal biome. Phylogenomic reconstruction based on 120 conserved bacterial marker genes, together with Average Nucleotide Identity (ANI >97.6%) and digital DNA–DNA hybridization (dDDH 77.7–90.4%) analyses, confirmed their assignment to *M. faecalis* and demonstrated high genomic relatedness to strains previously recovered from soil and human feces samples. Pangenome analysis of five available genomes revealed a total repertoire of 9,827 genes, with a large core genome comprising 7,499 genes (76.3%), consistent with a highly conserved and nearly closed pangenome structure. Functional classification based on COG categories showed uniform distributions across all isolates. Comparative analysis of the degradome further revealed strong conservation of proteolytic and carbohydrate-active enzyme repertoires, dominated by serine and metallopeptidases and diverse glycoside hydrolases. The extensive genomic and functional similarity among isolates from geographically distant and ecologically distinct environments supports a broad ecological distribution of *M. faecalis* and suggests that its large and conserved genomic repertoire underpins its persistence across contrasting habitats. These findings expand the known ecological range of the species and provide a comparative genomic framework for future investigations into its distribution and functional potential across different habitats.

## 1. Introduction

The phylum Myxococcota represents a group of social bacteria renowned for their complex life cycles and predatory behavior (Li et al., 2024; Waite et al., 2020). Broadly referred to as myxobacteria, these microorganisms are typically found in terrestrial habitats, such as soil, decaying wood and rodent dung, and in coastal and marine environments, either free-living or in association to other organisms (Saggu et al., 2023). The genus *Myxococcus* stands as the historical cornerstone of myxobacterial research. Due to a robust legacy of physiological and genomic studies, species like *M. xanthus* continue to serve as the principal models for deciphering the intricate life cycles and social interactions that define these microorganisms (Contreras-Moreno et al., 2024; Lall et al., 2024; Zhang et al., 2021).

The species *M. faecalis* was described from isolates recovered from human fecal samples (Das et al., 2025a) and is presently represented in the GenBank database by three genomes: strains O35 and O15, recovered from fecal samples obtained from Indian patients, and strain KYC1117, isolated from soil in South Korea (Das et al., 2025a; Park et al., 2025). Notably, although the species was formally described from clinical specimens, metagenomic analyses failed to detect the lineage in human-associated microbiomes, identifying it instead in soil and rhizosphere datasets (Das et al., 2025a). This discrepancy suggests that *M. faecalis* is a cosmopolitan environmental generalist with an ecological distribution that remains largely undocumented.

This ecological narrative is expanded herein through the exploration of mangrove ecosystems, which are unique intertidal zones characterized by high organic matter turnover and diverse microbial communities, including the complex social populations of myxobacteria (Octaviana et al., 2023; Zou et al., 2024). The present report describes the isolation and genomic characterization of *M. faecalis* strains BRX-014 and BRX-032 from mangroves on the coast of São Paulo State, in Southeastern Brazil, marking this as the first report of the species in a marine-coastal biome. While the strains were recovered from water or sediment samples collected nearly 100 km apart, both exhibit characteristic gliding motility and develop pigmented fruiting bodies in harsh nutritional settings. The portrayal of environmental *M. faecali*s isolates provides the first opportunity to evaluate the genetic repertoire of this species in a non-clinical, tropical ecosystem.

## 2. Material and Methods

### 2.1. Isolation of Strains

Strains BRX-014 and BRX-032 were isolated from coastal estuaries across the State of São Paulo (Southeastern Brazil) from samples obtained as part of a research project focused on assessing the myxobacterial diversity of tropical intertidal ecosystems. Collections were conducted under licenses from SISBIO no. 92515-2, SEMIL no. 000364/2024-87, and SISGEN A9D125D and AF72C6F. BRX-014 was recovered from water sampled from the banks of the Mar Casado Channel at the Portinho Mangrove, in the municipality of Praia Grande (23.986465° S, 46.403442° W). About 2 L of water was sieved against a filter-paper membrane, then a section of the membrane was plated onto ST21 solid media (see Table S1 for media composition). BRX-032, in turn, was isolated from sediments collected at the Barra do Sahy Mangrove, in the municipality of São Sebastião (23.774918° S, 45.690890° W). Sediment samples were placed onto WCX agar plates (see Table S1 for media composition) supplemented with heat-killed *Escherichia coli* inoculum, as baits. Inoculated plates were incubated at 30 °C and monitored daily for the development of swarms and fruiting bodies.

### 2.2. Whole Genome Sequencing, Assembly, and Annotation

The DNA from strains BRX-014 and BRX-032 was extracted with the DNeasy PowerSoil Pro Kit, quantified and evaluated for purity with NanoDrop™, and assessed for integrity by 1% gel electrophoresis. The sample was then sent to Instituto de Pesquisa do Câncer (IPEC; Brazil) for sequencing using the NovaSeq 6000 platform. The raw sequencing reads were quality-controlled and trimmed using Trimmomatic (version 0.39) to remove low-quality bases and adapter sequences (Bolger et al., 2014). The clean reads were then assembled into contigs using SPADES (version 3.13.1) with k-mer sizes of 77, 87, 99, and 121 (Bankevich et al., 2012). Quality of the genome assembly was assessed using CheckM (v1.0.18), as implemented in the KBase platform, which estimated genome completeness and contamination based on the presence of lineage-specific marker genes (Arkin et al., 2018; Parks et al., 2015). This step ensured that the assembly contained a high proportion of conserved genes, indicating good coverage and completeness. For genome annotation, the assembled genome was annotated using PROKKA (version 1.13) (Seemann, 2014).

### 2.3. Comparative Genomic Studies and Taxonomic Affiliation

To confirm the taxonomic classification and assess the genetic relatedness of strains BRX-014 and BRX-032, pairwise genomic comparisons were performed against *M. faecalis* reference strains (KYC1117, O15, and O35) sourced from GenBank. Average Nucleotide Identity (ANI) values were calculated using FastANI (v1.33) (Jain et al., 2018). A cut-off value of ≥95% was considered for species delineation. Additionally, digital DNA-DNA Hybridization (dDDH) values were estimated using the Genome-to-Genome Distance Calculator (GGDC 3.0) web server (http://ggdc.dsmz.de) (Meier-Kolthoff et al., 2013). Formula 2 (identities / HSP length) was selected for the analysis as it is robust against the effects of using incomplete draft genomes.

### 2.4. Pangenome and Phylogenomic Analysis

To reconstruct the pangenome of the *M. faecalis* species, the annotated GFF3 files were processed using Panaroo (v1.2.9) (Tonkin-Hill et al., 2020) in strict mode to cluster orthologous gene families and analyze the distribution of core and accessory genes across isolates. Taxonomic classification and phylogenomic reconstruction were performed using the GTDB-Tk toolkit (v2.3.0) (Chaumeil et al., 2022). The classify_wf workflow was employed to identify and align a set of 120 single-copy bacterial marker genes (bac120). The resulting concatenated multiple sequence alignment (MSA) was used to infer the phylogenetic tree and confirm the taxonomic placement of strains BRX-014 and BRX-032 within the *M. faecalis* clade using the Genome Taxonomy Database (GTDB) reference data.

### 2.5. Functional Annotation and Degradome Profiling

The functional potential of the pangenome was categorized using the COG (Clusters of Orthologous Groups) database via the eggNOG-mapper (v2.1.9) pipeline (Cantalapiedra et al., 2021). To specifically characterize the proteolytic and carbohydrate-degrading repertoire, specialized databases were employed. Peptidase-encoding sequences were identified by a local BLASTp search against the MEROPS database (Release 12.4) (Rawlings et al., 2009). Carbohydrate-Active Enzymes (CAZymes) were annotated using the dbCAN3 standalone tool (Zheng et al., 2023). The subcellular localization of all identified proteases and CAZymes was predicted using DeepLoc 2.0 (Thumuluri et al., 2022) determine their potential secretion.

## 3. Results

### 3.1. Genomic Features and Phylogenomic Placement

The genomic features of the mangrove isolates were highly congruent with those of the *M. faecalis* reference genomes (Tables 1 and S2). The genome sizes of BRX-014 (10.86 Mb) and BRX-032 (10.40Mb) were within the range observed for the clinical references O35 (10.53 Mb) and O15 (10.99 Mb), and the soil-derived strain KYC1117 (10.77 Mb). Furthermore, G+C content was identical across all five isolates (70%). The taxonomic identity of strains BRX-014 and BRX-032 was confirmed through a polyphasic approach combining 16S rRNA gene phylogeny and phylogenomic analysis. The 16S rRNA gene-based phylogeny placed both Brazilian isolates within the genus *Myxococcus*, specifically clustering them within the *M. faecalis* clade, sharing high sequence identity (99.8%) with the type strain *M. faecalis* (O35) (Fig. S1). Furthermore, a phylogenomic reconstruction utilizing 120 universal marker genes corroborated the classification of these Brazilian mangrove isolates as *M. faecalis*. These strains were nested within a robust monophyletic clade alongside reference strains from India (clinical) and South Korea (soil) (Fig. S2).

**Table 1.**
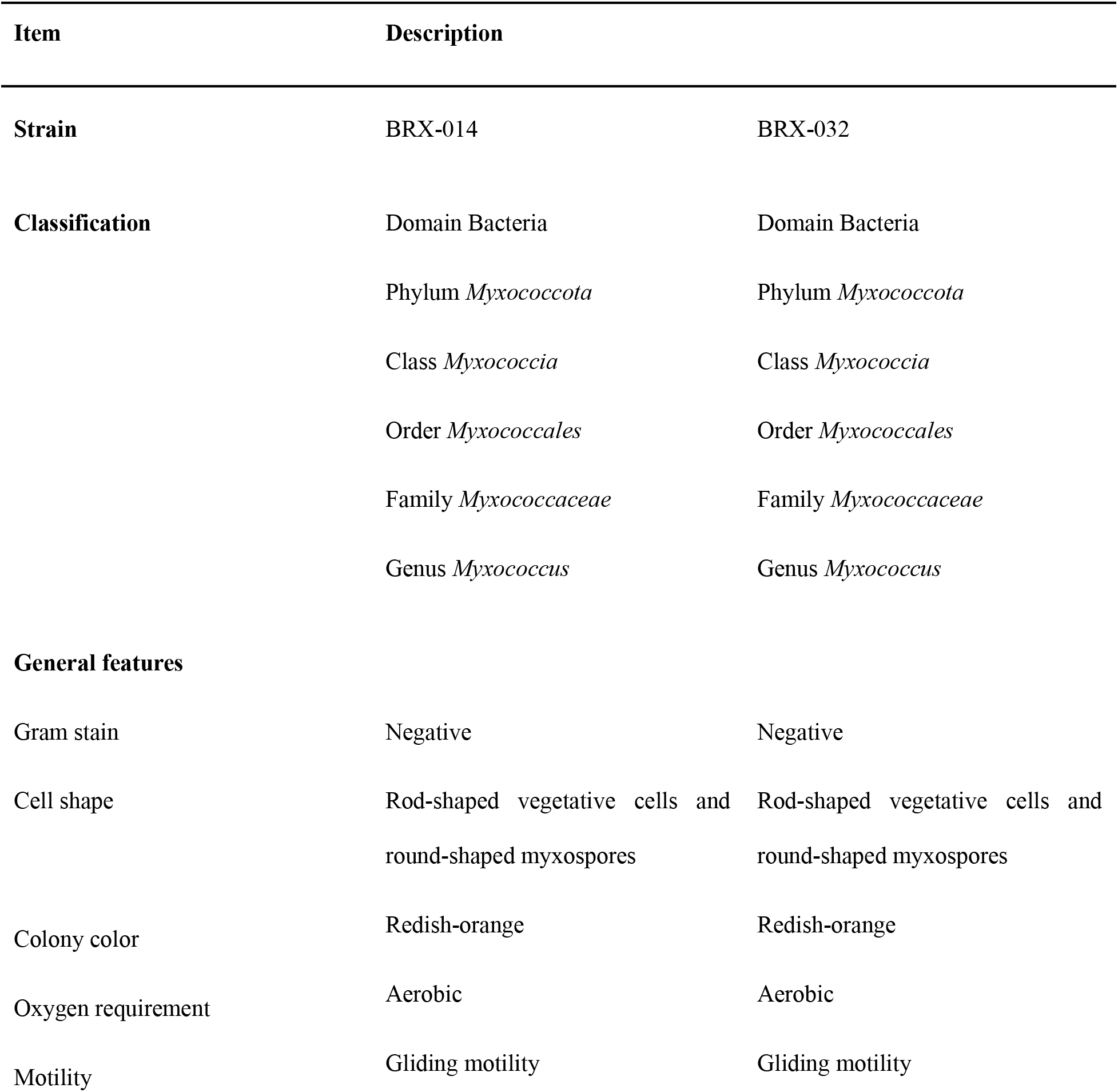
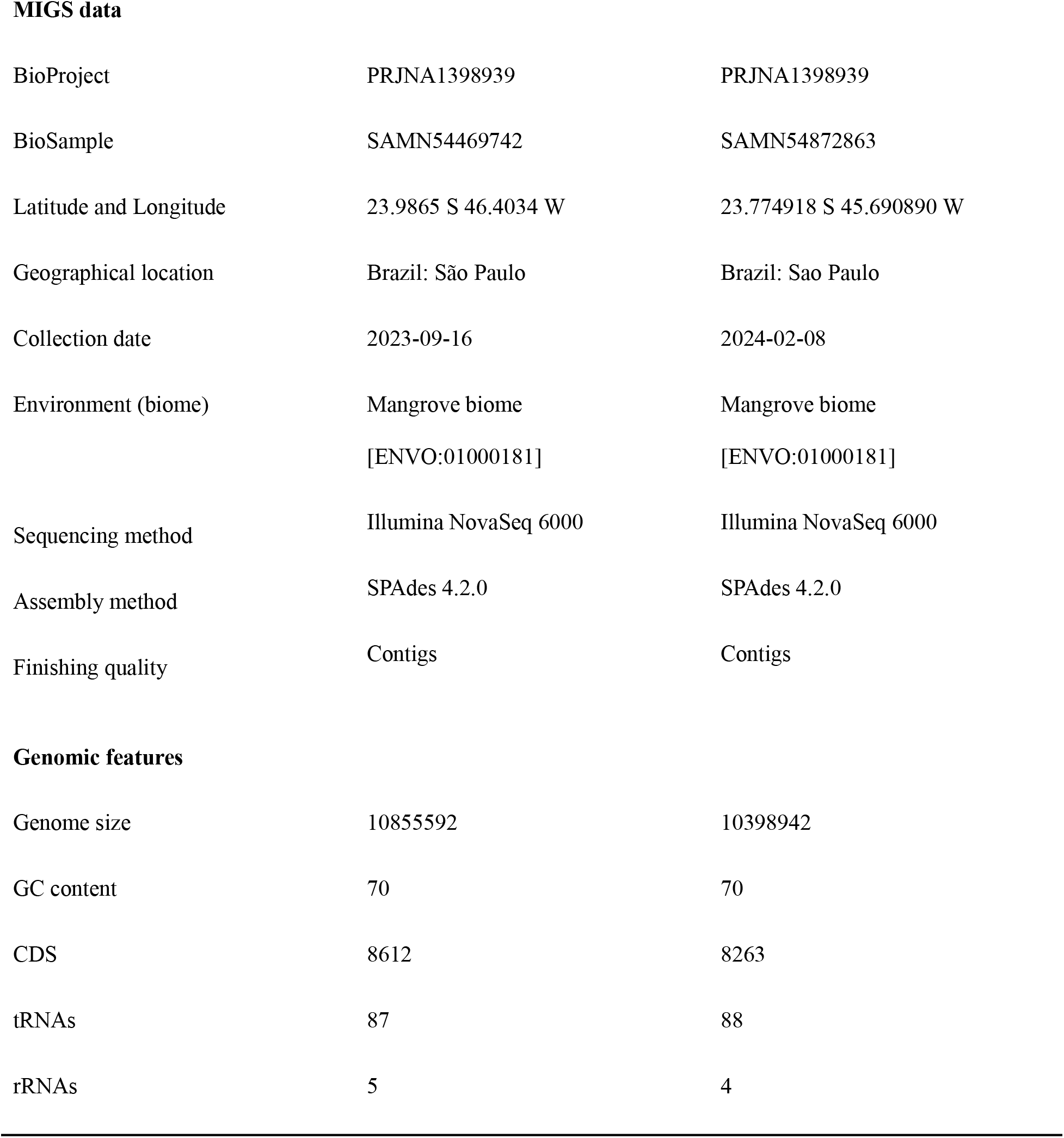
General features of *Myxococcus faecalis* BRX-014 and BRX-032.

| Item | Description |  |
| --- | --- | --- |
| Strain | BRX-014 | BRX-032 |
| Classification | Domain Bacteria | Domain Bacteria |
|  | Phylum *Myxococcota* | Phylum *Myxococcota* |
|  | Class *Myxococcia* | Class *Myxococcia* |
|  | Order *Myxococcales* | Order *Myxococcales* |
|  | Family *Myxococcaceae* | Family *Myxococcaceae* |
|  | Genus *Myxococcus* | Genus *Myxococcus* |
| General features |  |  |
| Gram stain | Negative | Negative |
| Cell shape | Rod-shaped vegetative cells and round-shaped myxospores | Rod-shaped vegetative cells and round-shaped myxospores |
| Colony color | Redish-orange | Redish-orange |
| Oxygen requirement | Aerobic | Aerobic |
| Motility | Gliding motility | Gliding motility |
| **MIGS data** |  |  |
| BioProject | PRJNA1398939 | PRJNA1398939 |
| BioSample | SAMN54469742 | SAMN54872863 |
| Latitude and Longitude | 23.9865 S 46.4034 W | 23.774918 S 45.690890 W |
| Geographical location | Brazil: São Paulo | Brazil: Sao Paulo |
| Collection date | 2023-09-16 | 2024-02-08 |
| Environment (biome) | Mangrove biome<br>[ENVO:01000181] | Mangrove biome<br>[ENVO:01000181] |
| Sequencing method | Illumina NovaSeq 6000 | Illumina NovaSeq 6000 |
| Assembly method | SPAdes 4.2.0 | SPAdes 4.2.0 |
| Finishing quality | Contigs | Contigs |
| **Genomic features** |  |  |
| Genome size | 10855592 | 10398942 |
| GC content | 70 | 70 |
| CDS | 8612 | 8263 |
| tRNAs | 87 | 88 |
| rRNAs | 5 | 4 |

To achieve definitive species-level resolution, Overall Genome Relatedness Indices (OGRI) were calculated against the three *M. faecalis* genomes currently available in GenBank: the Indian clinical strains (O35 and O15) and the soil-derived isolate KYC1117 from South Korea. Marine-derived strains BRX-014 and BRX-032 exhibited Average Nucleotide Identity (ANI) values exceeding 97.6% and digital DNA-DNA hybridization (dDDH - Formula 2) values ranging from 77.7% to 90.4% when compared to both clinical and terrestrial references (Fig. 1). These results significantly surpass the established thresholds for bacterial species delimitation (95–96% for ANI and 70% for dDDH), confirming the taxonomic classification and establishing a high degree of genomic conservation across the species, regardless of the geographic or environmental origin of the isolates (Fig. 2).

**Figure 1.**
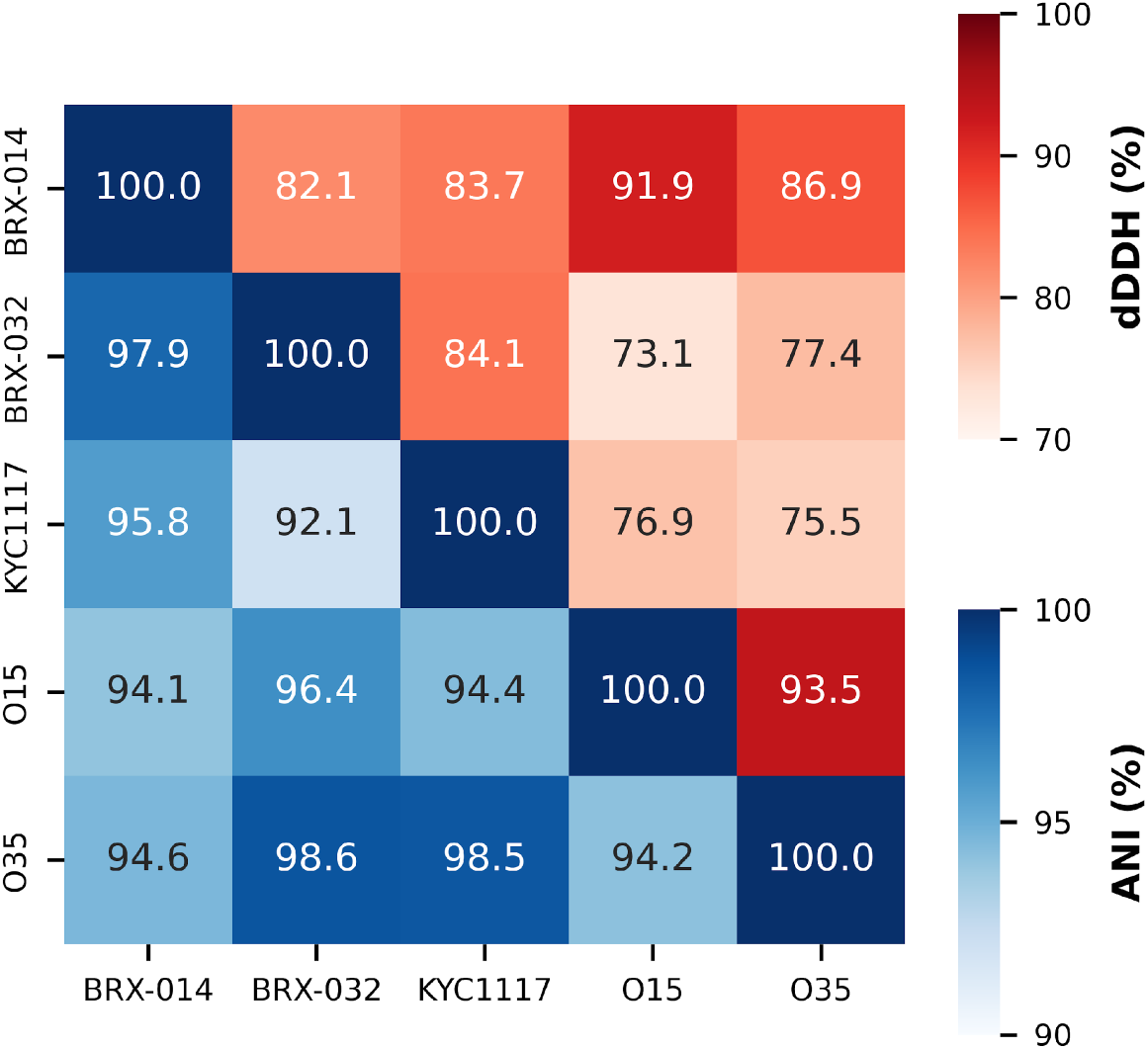
Comparative genomic metrics of *Myxococcus faecalis* strains. Heatmap displaying pairwise Average Nucleotide Identity (ANI) values in the lower triangle (blue scale) and digital DNA-DNA Hybridization (dDDH) values in the upper triangle (red scale) for strains BRX-014, BRX-032, KYC1117, O15, and O35.

**Figure 2.**
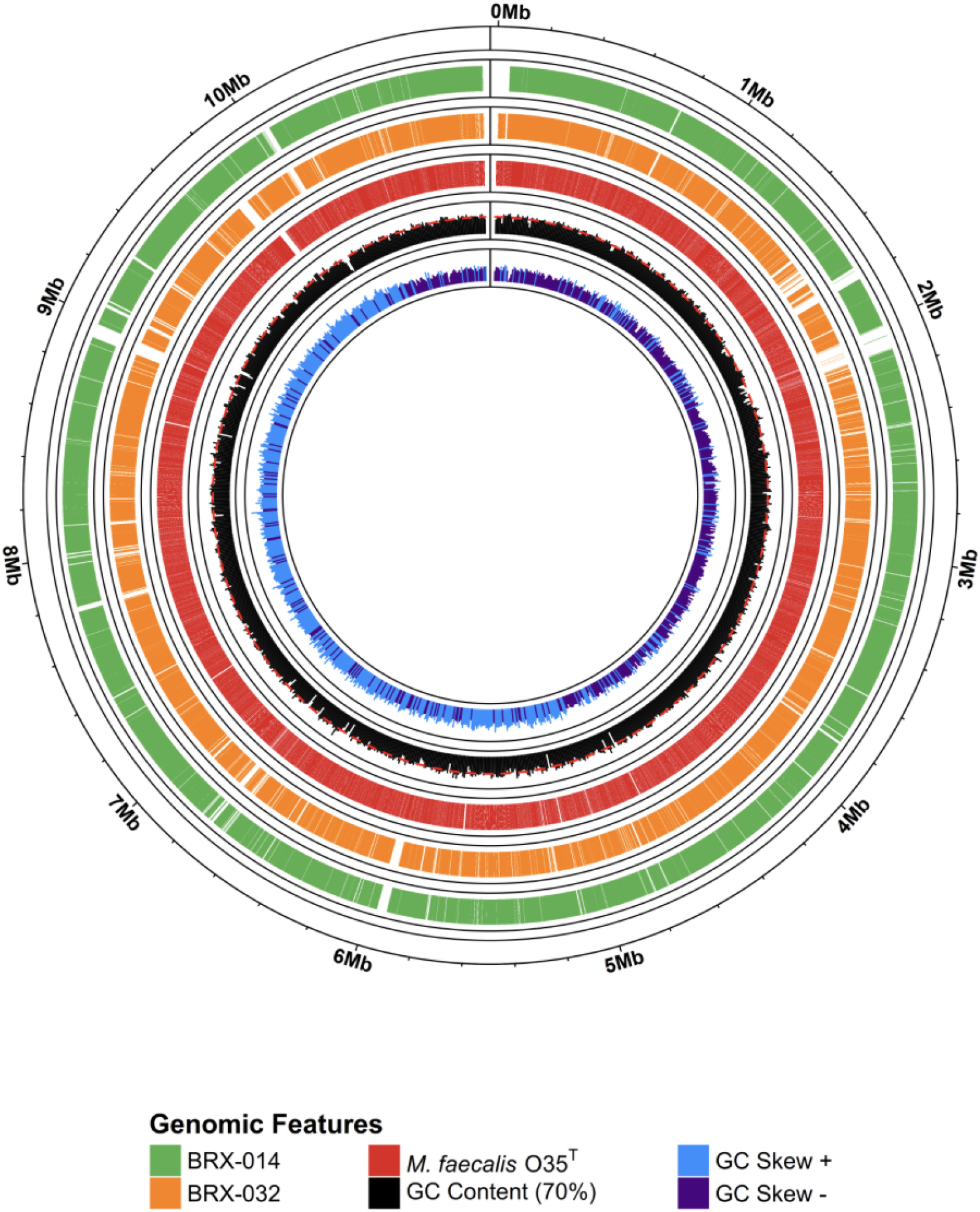
Circular genome map of *Myxococcus faecalis* O35 and comparative analysis with mangrove isolates. From the outer to the inner tracks: (1) genomic coordinates in Megabases (Mb); (2) alignment with isolate BRX-014 (green); (3) alignment with isolate BRX-032 (orange); (4) Coding Sequences (CDS) of the type strain (red); (5) GC content (black) with a 70% baseline; and (6) GC skew (positive in blue and negative in purple).

### 3.2. Comparative Pangenome Analysis

To explore the genomic diversity and evolutionary stability of *M. faecalis*, a pangenome analysis was performed comparing the two Brazilian mangrove isolates with the three publicly available genomes. The total pangenome of the species comprises 9,827 genes, characterized by a remarkably high degree of conservation. The core genome (present in all five strains) consists of 7,499 genes, representing approximately 76.3% of the total pangenome repertoire. The presence of a substantial core genome highlights a high level of genomic conservation, representing a functional repertoire that remains stable across disparate geographical regions and ecological niches. No “soft core” or “cloud” genes were identified in this set of five genomes, indicating that the genetic variation is strictly confined to the shell genome (2,328 genes, ~23.7%). The Flower Plot (Fig. 3A) depicts the distribution of gene clusters, with the core genome at the center and shared gene clusters represented by the intersecting regions. Notably, although unique genes are present within the shell fraction, they account for only a minor proportion of the total genetic repertoire.

**Figure 3.**
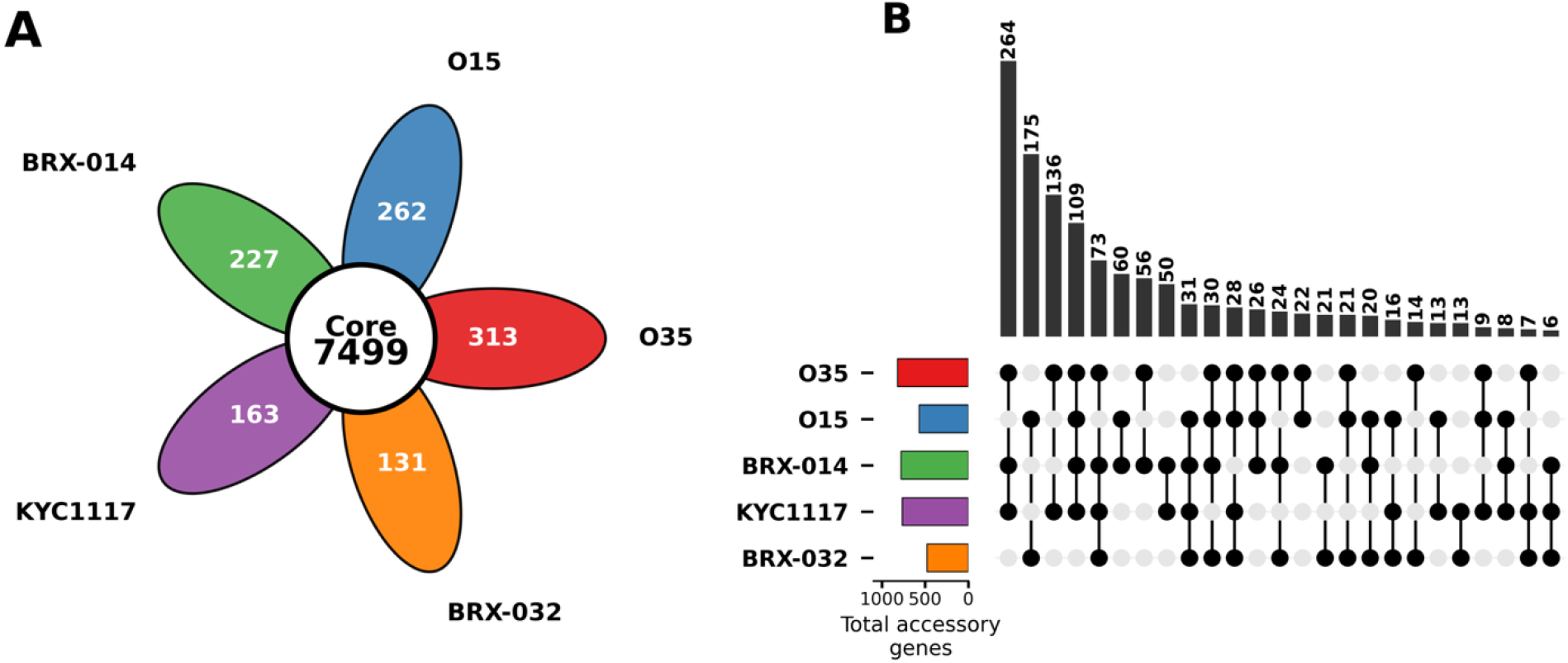
Pangenome and accessory genome intersections of *Myxococcus faecalis* strains. **(A)** Flower plot illustrating the core genome (center) and unique genes (petals) for five strains. **(B)** UpSet plot showing accessory genome intersections; vertical bars represent intersection size, while the matrix indicates strain membership. The bottom-left bar chart shows the total accessory gene repertoire per strain. Brazilian mangrove isolates are color-coded: BRX-014 (green) and BRX-032 (orange).

Beyond the core set, the accessory genome showcased distinct patterns of gene sharing. Notably, the largest accessory intersection (n = 264 genes) was shared exclusively between the mangrove isolate BRX-014, the soil strain KYC1117, and the clinical strain O35. In contrast, BRX-032 exhibited its primary accessory affiliation with the clinical strain O15, sharing 175 genes. The unique gene content (singletons) also varied significantly among isolates (Fig. 3B). The distribution of the accessory genome closely mirrors the evolutionary relationships observed in the phylogenomic reconstruction, reinforcing the high degree of consistency between lineage descent and gene content. Strains that clustered together in the marker-gene-based tree also exhibited a higher degree of shared genomic repertoire in the pangenome analysis, corroborating the evolutionary stability of these lineages (Fig. 1, S1, and S2). This structure suggests a “closed” or “nearly closed” pangenome for *M. faecalis*, where the addition of new genomes from geographically distant and ecologically distinct environments, such as the tropical mangroves of the South Atlantic, does not significantly expand the overall gene pool of the species.

### 3.3. Functional Classification of the Pangenome

The total genomic repertoire of the five *M. faecalis* strains was classified into functional categories using the COG database. The distribution of functional classes was remarkably uniform across all isolates, regardless of their origin (Fig. 4). The most prevalent category in all genomes was [S] Function Unknown (~20.6–20.9%), followed by [K] Transcription (~8.6–8.8%) and [T] Signal Transduction Mechanisms (~8.6–8.7%). These results indicate that while phylogenetic branching and accessory gene differences (as seen in the Fig. 3B) drive the fine-scale diversification of these strains, the fundamental functional framework of *M. faecalis* remains strictly conserved across diverse ecological niches.

**Figure 4.**
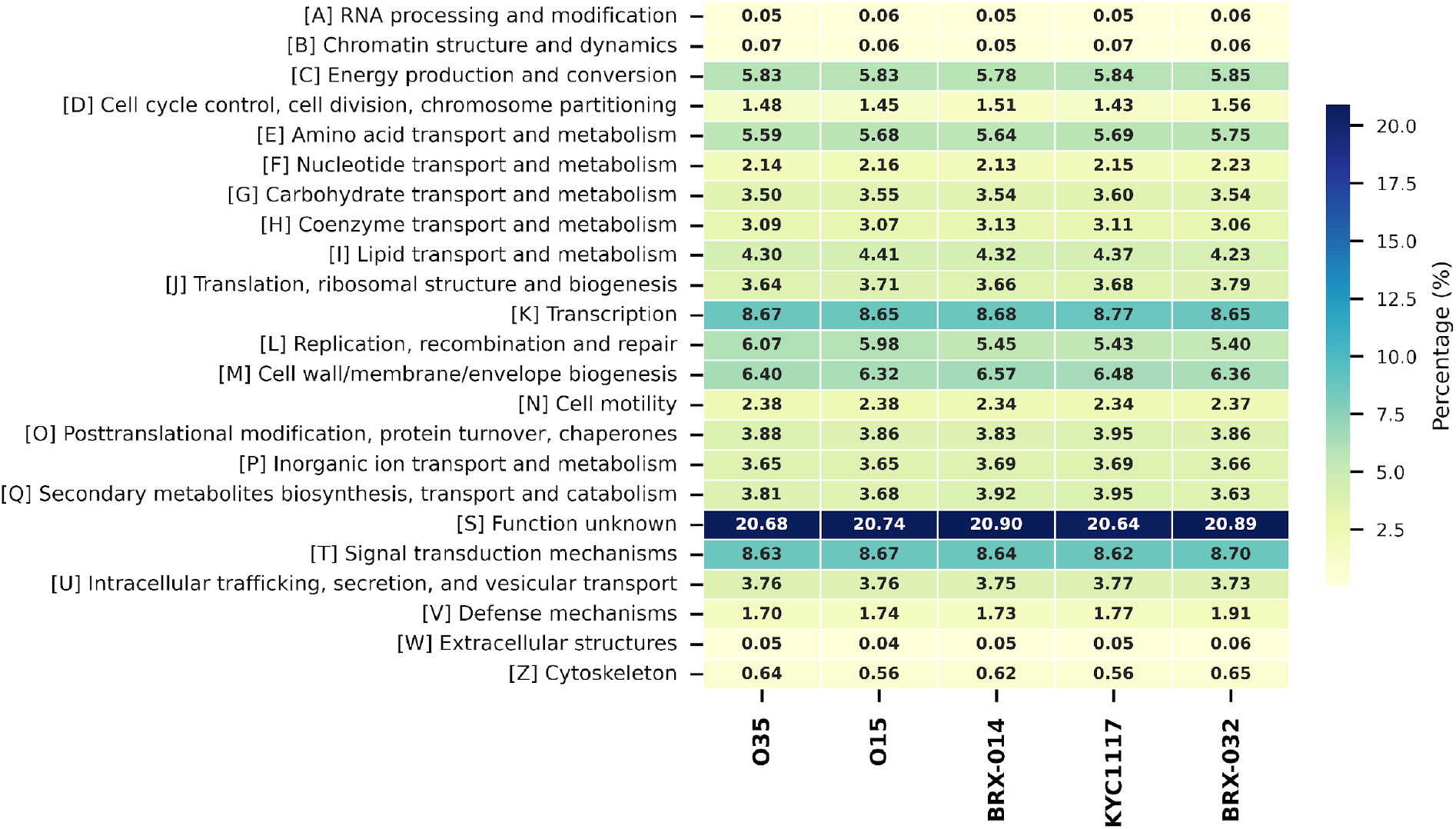
Functional profile of individual *Myxococcus faecalis* genomes based on COG categories. The heatmap displays the percentage distribution of genes across Clusters of Orthologous Groups (COG) for each of the five studied strains. Color intensity and numerical values represent the relative abundance of genes within each category per genome.

The proteolytic repertoire of the *M. faecalis* isolates was characterized by identifying peptidase-encoding sequences and their respective catalytic classes through the MEROPS database. Like the functional categories, distribution of these enzymes was remarkably consistent across all five genomes. The genomic distribution was predominantly composed of two catalytic groups: serine peptidases, which accounted for 41.76% to 42.46% of the identified sequences, and metallopeptidases, comprising 35.27% to 36.29%. Collectively, these two classes represent over 77% of the total peptidase repertoire in all analyzed isolates. Cysteine peptidases constituted the third most abundant group, with a stable occurrence between 9.72% and 10.82%. Sequences assigned to peptidases of unknown catalytic type represent approximately 6.01% to 6.55% of the total hits (Fig. S3). This high degree of uniformity in the proteolytic genomic content across geographically distant biomes highlights a stable functional core in *M. faecalis*.

The carbohydrate-metabolizing potential of the *M. faecalis* strains was explored by identifying Carbohydrate-Active Enzymes (CAZymes) using the dbCAN3 pipeline. Like the proteolytic profile, the CAZyme repertoire exhibited a high degree of conservation across all five genomes, reflecting a specialized machinery for complex carbohydrate degradation. The most abundant classes identified were Glycoside Hydrolases (GH) and Glycosyltransferases (GT) which, together, constitute the core of the carbohydrate-active degradome. Auxiliary Activity (AA) enzymes and Carbohydrate Esterases (CE) were also present in consistent proportions, while Polysaccharide Lyases (PL) and Carbohydrate-Binding Modules (CBM) represented smaller fractions of the total repertoire. Analysis of subcellular localization via DeepLoc 2.0 revealed that, while a significant portion of the CAZyme machinery is predicted to be cytoplasmic, particularly GTs involved in cell wall biosynthesis and internal glycan processing, a substantial diversity of GHs was localized in the extracellular space and periplasm (Fig. S3).

This extracellular prevalence of GHs in this species suggests a robust capability for the breakdown of environmental polysaccharides. The presence of these enzymes in the secretome combined with CBM-containing proteins for substrate anchoring underscores the scavenger lifestyle of *M. faecalis* and its potential for biomass degradation in diverse marine and terrestrial ecosystems.

## 4. Discussion

Mangroves are unique intertidal ecosystems that provide specialized niches for a high abundance and diversity of microorganisms, including members of the phylum Myxococcota (Octaviana et al., 2023; Zou et al., 2024), which account for approximately 1.56% of the prokaryotic community in these saline sediments (Zou et al., 2024). These sophisticated social micro-predators function as key ecological connectors within the mangrove microbial community (Zou et al., 2024) and represent a promising reservoir for the discovery of novel specialized metabolites and antimicrobial compounds (Octaviana et al., 2023).

The genomic characterization of myxobacteria strains BRX-014 and BRX-032 provides intriguing insights into the ecology and global distribution of *M. faecalis*. The isolation of this species from Brazilian mangroves, together with previous reports from South Korean soil (KYC1117) and human fecal samples in India (O35 and O15), highlights a remarkably cosmopolitan distribution (Das et al., 2025b; Park et al., 2025). Despite vast geographical distances and stark physicochemical contrasts among these habitats, comparative genomics revealed an exceptionally high degree of conservation. The striking similarity in both genomic content and functional profiles (COG categories) across clinical, terrestrial, and marine isolates raises fundamental questions about the ecological role of *M. faecalis* in these diverse biomes. The enzymatic repertoire also displayed a notable level of conservation, with comparable distributions of catalytic classes and carbohydrate-active enzymes among all strains, further supporting a shared metabolic framework and functional stability across environments.

While myxobacteria are renowned for their metabolic versatility, the lack of significant functional differentiation in the mangrove isolates suggests that these populations are not undergoing substantial gene acquisition for local adaptation. This apparent genomic stasis is supported by the “microbial seed bank” hypothesis (Lennon and Jones, 2011; Shoemaker and Lennon, 2018), whereby bacteria capable of entering deep dormancy can persist in diverse environments, uncoupled from immediate environmental selection. Myxobacteria are well known for their complex developmental cycle, culminating in the formation of stress-resistant myxospores (Mohr, 2018). The production of such defiant structures is a critical mechanism for persistence against environmental fluctuations, including UV radiation and desiccation (Lall et al., 2024).

While this trait facilitates passive dispersal, evolutionary outcomes differ markedly between species. In the model organism *M. xanthus*, limited dispersal leads to a clear pattern of isolation by distance, where genetic differentiation increases significantly with geographic separation (Vos and Velicer, 2006). In contrast, the comparative analysis carried out herein of these five *M. faecalis* strains suggests a lack of such geographic structuring. The extensive genomic identity and uniform functional repertoire retained between South Atlantic and Asian isolates indicate that *M. faecalis* populations are likely not diverging via local adaptation. This genomic stasis supports the hypothesis that the species is maintained by a global “seed bank,” where dormancy uncouples the organism from local selective pressures, preserving species cohesion across disparate biomes (Shoemaker and Lennon, 2018).

In this scenario, the recovery of closely related genotypes from Brazilian mangroves and Asian soils likely reflects the ubiquitous distribution of *M. faecalis* spores. This also suggests that its presence in human fecal samples may be transient, representing ingested environmental spores rather than a true symbiotic or pathogenic relationship (David et al., 2014). This interpretation is consistent with the original species description, which noted that despite successful cultivation from clinical specimens, *M*. faecalis is absent from extensive human gut metagenomic datasets (Das et al., 2025a).

An alternative, non-mutually exclusive explanation for the observed cosmopolitan distribution of *M. faecalis* is its inherent ecological plasticity. Myxobacteria possess some of the largest genomes among prokaryotes, typically ranging from 9.0 to 16.0 Mbp (Pérez et al., 2022; Wang et al., 2023), which supports a generalist lifestyle and complex social behaviors (Muñoz-Dorado et al., 2016). Taking the strains analyzed herein, this is reflected in the high degree of genome conservation, with 7,499 core genes shared across marine, terrestrial, and clinical isolates. Such a large and conserved gene set likely allows vegetative cells to maintain metabolic versatility, enabling survival in multiple environments through differential gene expression rather than extensive genomic restructuring (Guieysse and Wuertz, 2012; Pérez et al., 2022; Wang et al., 2023).

This genomic strategy is an evolutionary response to variable and nutrient-limited ecological niches (Guieysse and Wuertz, 2012), allowing the predator to precisely adapt to different prey and environmental conditions by regulating its feeding and metabolic machinery (Pérez et al., 2022). However, while these mechanisms are increasingly understood in model species, the specific transcriptomic shifts and metabolic trade-offs that drive the global success of *M. faecalis* must be better studied to fully elucidate its unique ecological success Both the potential for long-term survival as spores and the metabolic flexibility of vegetative cells may depend on a substantial fraction of the *M. faecalis* genome that remains functionally uncharacterized. The conservation of these large genomic regions across isolates from contrasting habitats suggests that they may play a central role in the ecological success and resilience of this species, providing a genetic toolkit for survival across spatially and physicochemically diverse environments. Despite the insights gained from model myxobacteria, the specific contribution of these uncharacterized regions to the global distribution of *M. faecalis* remains unknown. Therefore, the functional role of these conserved hypothetical genes and their transcriptomic response to environmental fluctuations deserve to be better studied in *M. faecalis*. Elucidating these mechanisms will be essential to fully understand the ecological plasticity that allows this species to thrive across the diverse habitats of our planet.

## 5. Nucleotide sequence accession number

The complete genome sequences of *Myxococcus faecalis* strains BRX-014 and BRX-032 are deposited in the GenBank database under the accession numbers JBTMXL000000000 and JBTZSQ000000000, respectively, under project PRJNA1398939.

## CRediT authorship contribution statement

RSO: Conceptualization, Writing original draft, Review and editing, Software, Methodology, Data curation, Formal analysis. YFL: Writing original draft, Software, Methodology, Data curation, Formal analysis. PCJ: Conceptualization, Writing original draft, Review and editing, Supervision, Resources, Project administration.

## Acknowledgements

The authors wish to acknowledge the financial support from the following Brazilian agencies: São Paulo Research Foundation (FAPESP) in the form of grants 2023/08735-1 and 2022/12654-4; National Council for Scientific and Technological Development (CNPq) for grant 408549/2024-6; and Coordenação de Aperfeiçoamento de Pessoal de Nível Superior (CAPES) in the form of a doctoral scholarship to RSO (88887.969556/2024-00) and a masters scholarship to YFL (88887.283299/2026-00).

