## Supplementary_material for "Isolation and Genomic Characterization of *Myxococcus faecalis* Strains from Mangroves in Southeastern Brazil"

**Table S1. Detailed composition of culture media WCX and ST21.** Concentrations are expressed as percentage weight per volume (% w/v).

| Culture media | Composition % (w/v) | Reference |
|---|---|---|
| **WCX** | Agar: 1.4%<br>$CaCl_2 \cdot 2H_2O$: 0.1%<br>Cycloheximide: 0.0025% | Reichenbach, H., Dworkin, M. (1992). The Myxobacteria. In: Balows, A., Trüper, H.G., Dworkin, M., Harder, W., Schleifer, KH. (eds) The Prokaryotes. Springer, New York, NY. https://doi.org/10.1007/978-1-4757-2191-1_26 |
| **ST21** | Yeast extract: 0.002%<br>Agar: 1.4%<br>$K_2HPO_4$: 0.1%<br>$KNO_3$: 0.1%<br>$MgSO_4 \cdot 7H_2O$: 0.1%<br>$CaCl_2 \cdot 2H_2O$: 0.1%<br>$MnSO_4 \cdot 7H_2O$: 0.01%<br>$FeCl_3$: 0.1% | Reichenbach, H., Dworkin, M. (1992). The Myxobacteria. In: Balows, A., Trüper, H.G., Dworkin, M., Harder, W., Schleifer, KH. (eds) The Prokaryotes. Springer, New York, NY. https://doi.org/10.1007/978-1-4757-2191-1_26 |

**Table S2. Comparative genomic features of *Myxococcus faecalis* strains.** The table summarizes the assembly and quality metrics for the strains isolated in the current study (BRX-014 and BRX-032) and the reference genomes obtained from public databases (KYC1117, O35, and O15). Completeness and contamination [RO1] were estimated using CheckM2. Publicly available accession numbers: [1]GCF_046058965.1; [2]GCF_050039175.1; [3]GCF_050039385.1.

| Feature | BRX-014 | BRX-032 | KYC1117[1] | O35[2] | O15[3] |
|---|---|---|---|---|---|
| **Isolation Source** | Mangrove water | Mangrove sediment | Soil | Human feces | Human feces |
| **Genome Size (Mb)** | 10.86 | 10.40 | 10.77 | 10.53 | 10.99 |
| **G+C content (%)** | 70.0 | 70.0 | 70.0 | 70.0 | 70.0 |
| **Contig N50 (kb)** | 613.3 | 330.6 | 10,772.5 | 10,528.4 | 10,986.1 |
| **Total Contigs** | 58 | 79 | 1 | 1 | 1 |
| **Completeness (%)*** | 100.0 | 99.99 | 100.0 | 99.99 | 100.0 |
| **Contamination (%)*** | 01.03 | 01.09 | 0.99 | 1.22 | 1.66 |
| **Coding Density (%)** | 90.8 | 90.5 | 90.8 | 90.4 | 90.6 |
| **Total CDS** | 8,618 | 8,3 | 8,446 | 8,367 | 8,654 |

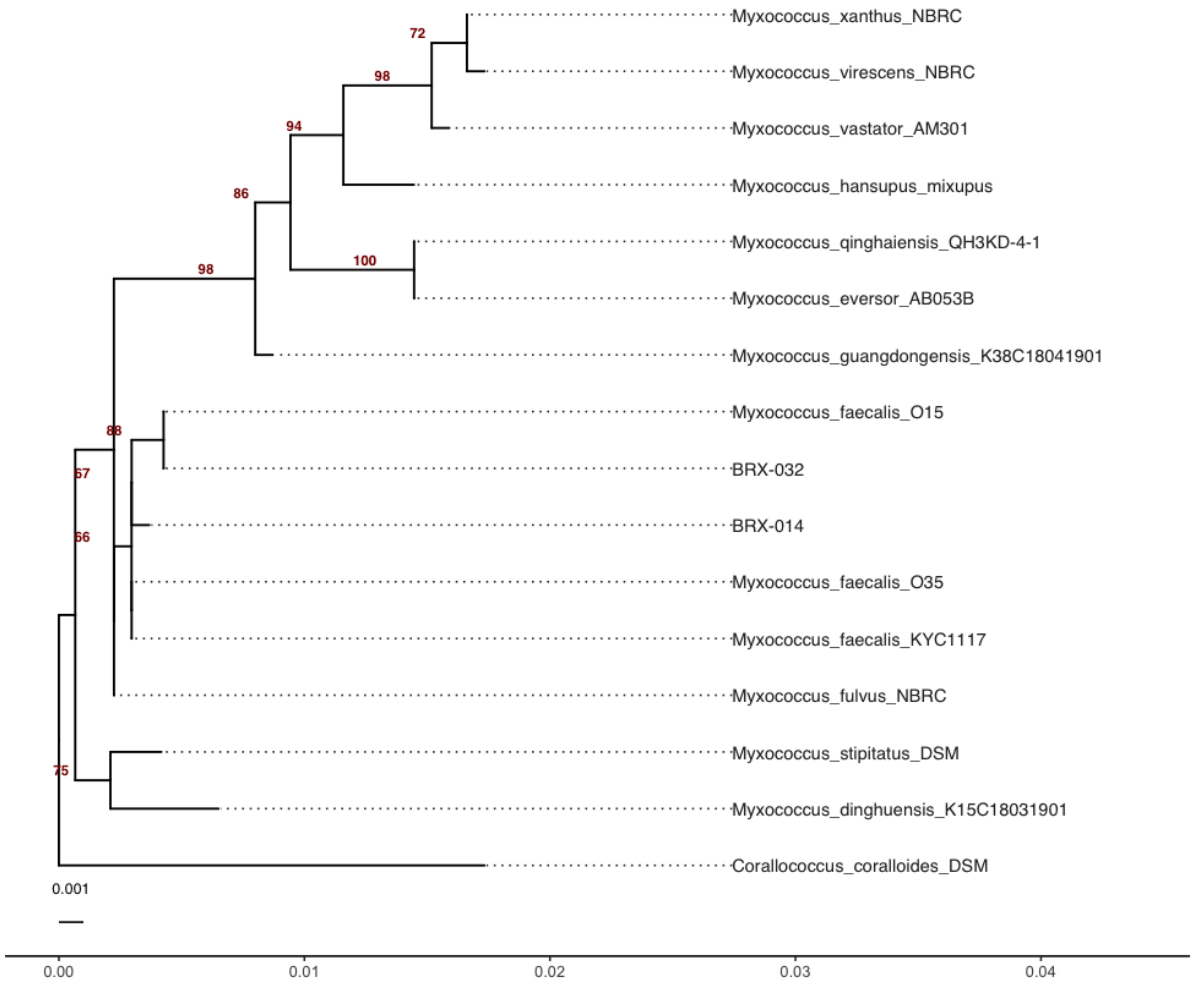


**Figure S1.** Phylogenetic tree based on 16S rRNA gene sequences. The tree was reconstructed using the Neighbor-Joining (NJ). Numerical values at the nodes represent bootstrap support percentages (from 1,000 replicates),

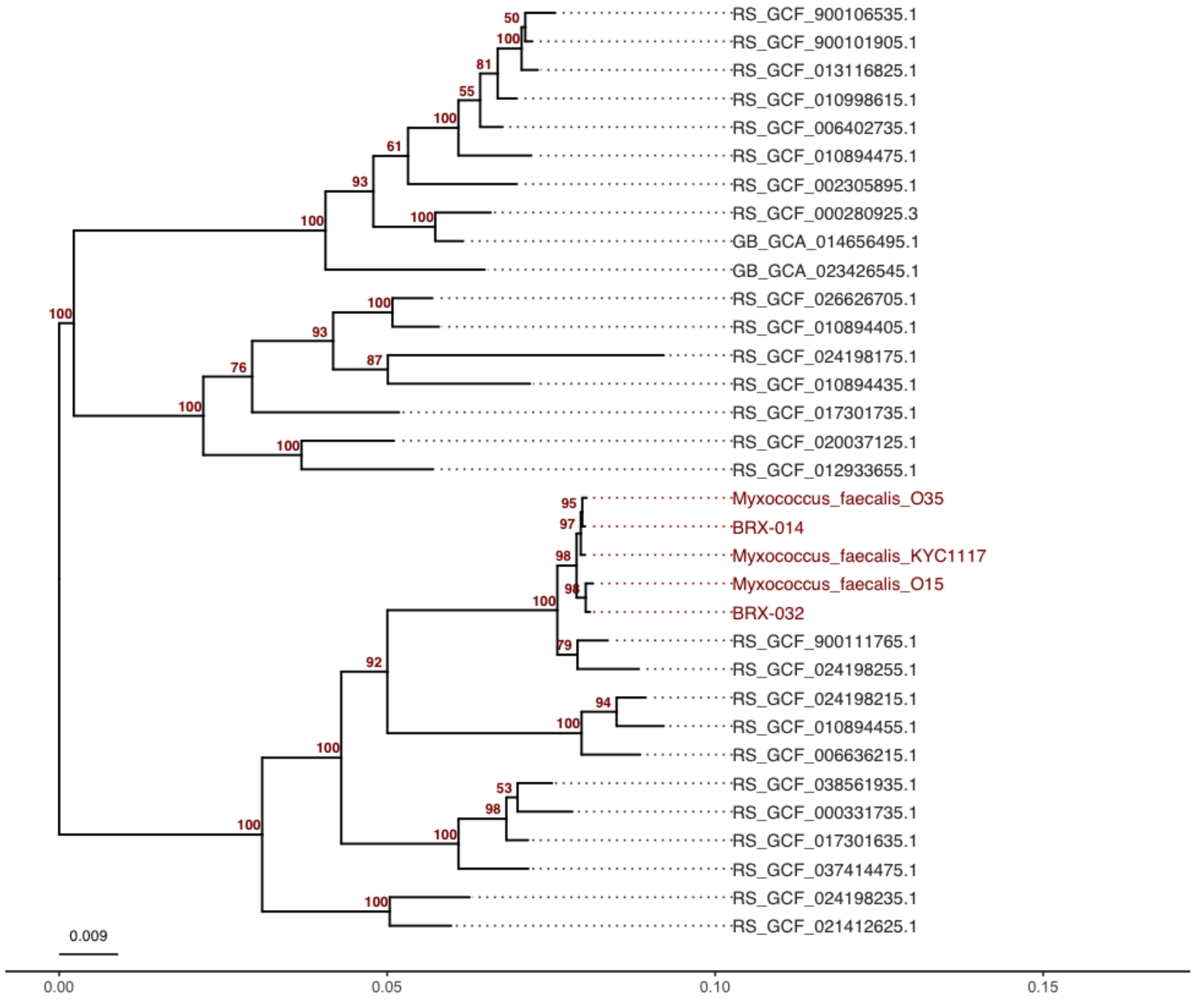


**Figure S2. Maximum-likelihood phylogenomic tree of *Myxococcus faecalis*.** Based on the analysis of 120 universal marker genes (GTDB), the Brazilian mangrove isolates cluster in a well-supported monophyletic group with reference strains from India and South Korea. Support values are indicated at the nodes.

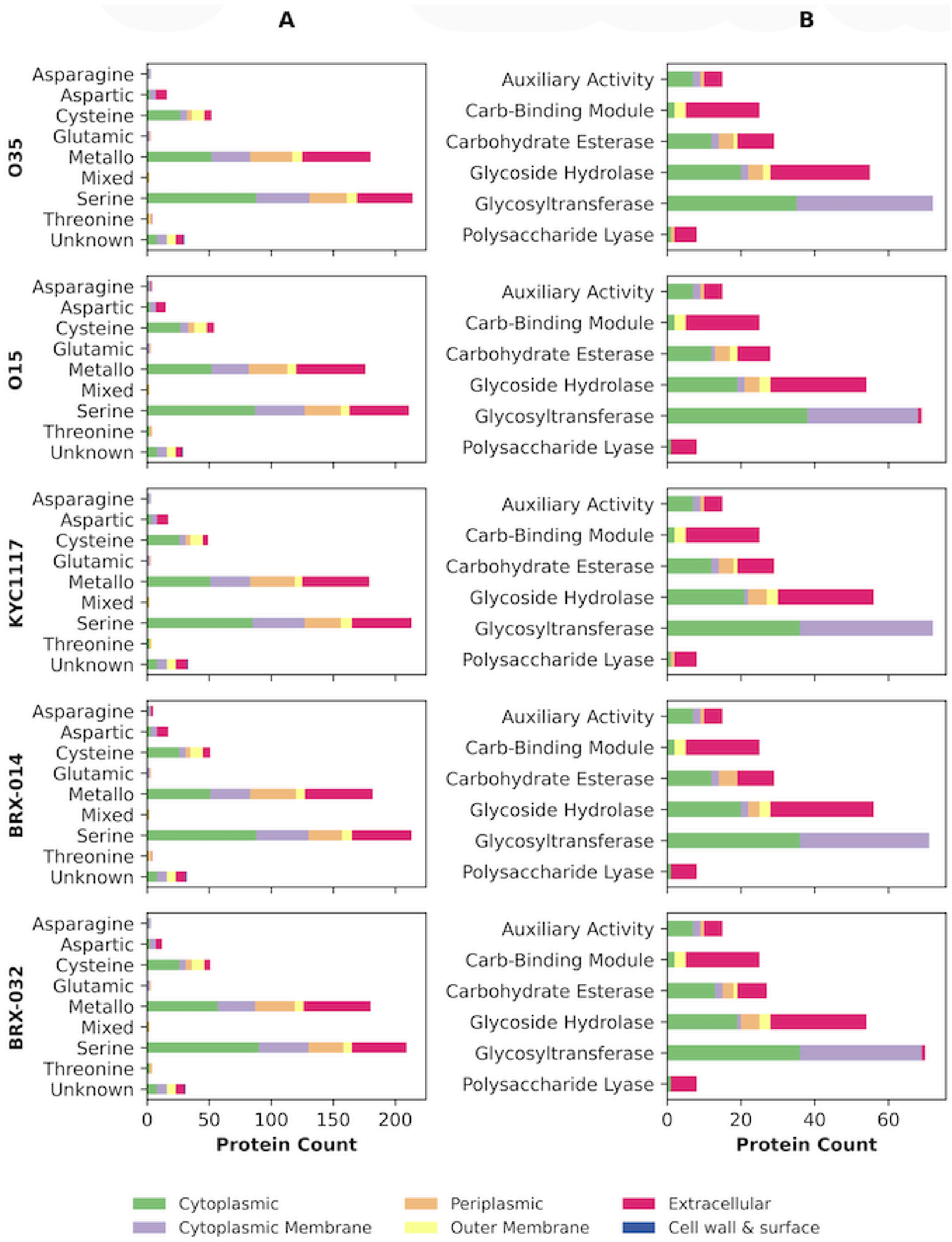


**Figure S3. Integrated degradome profiling and subcellular localization of *Myxococcus faecalis*.** Genomic abundance of protease classes (A) and CAZymes (B) for five *M. faecalis* isolates. Colors indicate the predicted subcellular localization for each enzyme class according to DeepLoc 2.0. Reference strains (O35, O15, KYC1117) and mangrove isolates (BRX-014, BRX-032) are organized by rows. Gene counts and functional classifications are based on MEROPS and dbCAN3annotations, respectively.
